# Spatial Dynamics and Emergent Properties of pLS20 Conjugation on Solid Surfaces

**DOI:** 10.1101/2025.02.27.640280

**Authors:** Álvaro López-Maroto, Wilfried J J Meijer, Javier Buceta, Saúl Ares

## Abstract

Horizontal gene transfer (HGT) is a major evolutionary process in bacteria, driving the dissemination of genetic traits including antibiotic resistance (AR). In this study, we employ a hybrid modeling approach, combining agent-based simulations and Ordinary Differential Equation (ODE) models, to investigate bacterial conjugation—a key HGT mechanism whose dynamics remain poorly understood. Our agent-based simulations of the transfer dynamics of the conjugative plasmid pLS20 from *Bacillus subtilis* reveal that spatial organization, colony growth dynamics, and quorum-sensing regulation significantly influence plasmid dissemination. Increased donor-recipient mixing enhances plasmid transmission by reducing quorum-induced repression, while colony growth-driven displacement of donor cells alters the local distribution of quorum-sensing signals, enabling sustained conjugation activity at the colony periphery. Complementary ODE modeling captures macroscopic trends in plasmid transmission, providing insights into the interplay between spatial factors and regulatory mechanisms. By bridging single-cell regulatory dynamics with population-level behaviors, this study advances our understanding of bacterial conjugation on solid surfaces, offering potential strategies for mitigating the spread of antibiotic resistance.

**Author summary:** Bacteria can exchange genetic material through a process called horizontal gene transfer, which helps them adapt to new environments but also develop traits like antibiotic resistance. One of the most important ways bacteria share genes is through conjugation—a mechanism where a conjugative element transfers from a donor to a recipient via a channel connecting both cells. Although considerable knowledge has been gathered over the last decades concerning regulation of the conjugation genes and the structure of the transferosome responsible for transfer of the conjugative element, far less is known about the dynamics of conjugative transfers within populations of cells, especially on solid medium.

Our study focuses on the conjugative plasmid pLS20 from *Bacillus subtilis*, a model bacterium related to several pathogens. Using computer simulations, we modeled how this plasmid spreads within bacterial colonies growing on solid surfaces. We found that the spatial organization of bacteria plays a large role: well-mixed populations allow the plasmid to spread more effectively, while certain growth patterns disrupt the signaling processes that regulate gene transfer. This study helps us better understand how bacteria transfer genes in complex environments and could guide new strategies to combat antibiotic resistance.

## Introduction

Horizontal gene transfer (HGT) is a fundamental biological process enabling the exchange of genetic material between organisms, facilitating rapid adaptation and evolution. HGT has been documented across all domains of life, including transfer among prokaryotes, between prokaryotes and eukaryotes, and even within eukaryotic lineages [1]. By bypassing traditional evolutionary paths, HGT drives genome innovation, impacting key processes such as antibiotic resistance (AR) acquisition and the dissemination of virulence factors [2]. The proliferation of AR, a major global health crisis, exemplifies the importance of understanding HGT mechanisms to devise strategies for mitigating its consequences [3–5]. Among the mechanisms of HGT, bacterial conjugation is particularly impactful, serving as a primary driver of AR and virulence gene dissemination [6].

Bacterial conjugation involves the transfer of genetic elements, such as plasmids or integrative elements, from a donor to a recipient cell via a direct physical connection. While conjugation is ubiquitous among both Gram-positive (G+) and Gram-negative (G-) bacteria, most studies have focused on G-species, leaving many aspects of G+ conjugation underexplored. This knowledge gap limits our ability to address AR dissemination in clinically significant G+ genera, such as *Listeria, Staphylococcus, Clostridium, Enterococcus*, and *Bacillus*. Furthermore, experimental challenges associated with studying bacterial conjugation at the population level hinder efforts to understand emergent properties that govern gene transfer dynamics in complex environments [3–5].

The process of bacterial conjugation can be divided into four distinct steps: initial cell-to-cell attachment, formation of a secretion channel, processing and transfer of the conjugative element, and establishment of the transferred genetic material in the recipient cell [7, 8]. These steps require a suite of specialized proteins, imposing a significant metabolic burden on donor cells. To mitigate this burden, conjugation systems are tightly regulated and remain in a default OFF state until environmental conditions favor activation [9].

In this study, we focus on the conjugative plasmid pLS20 from *Bacillus subtilis*, a well-characterized model organism closely related to several pathogenic G+ species [10–12]. The 65-kb pLS20 plasmid contains a large conjugation operon comprising all the conjugation genes (Fig 1A).

**Fig 1.**
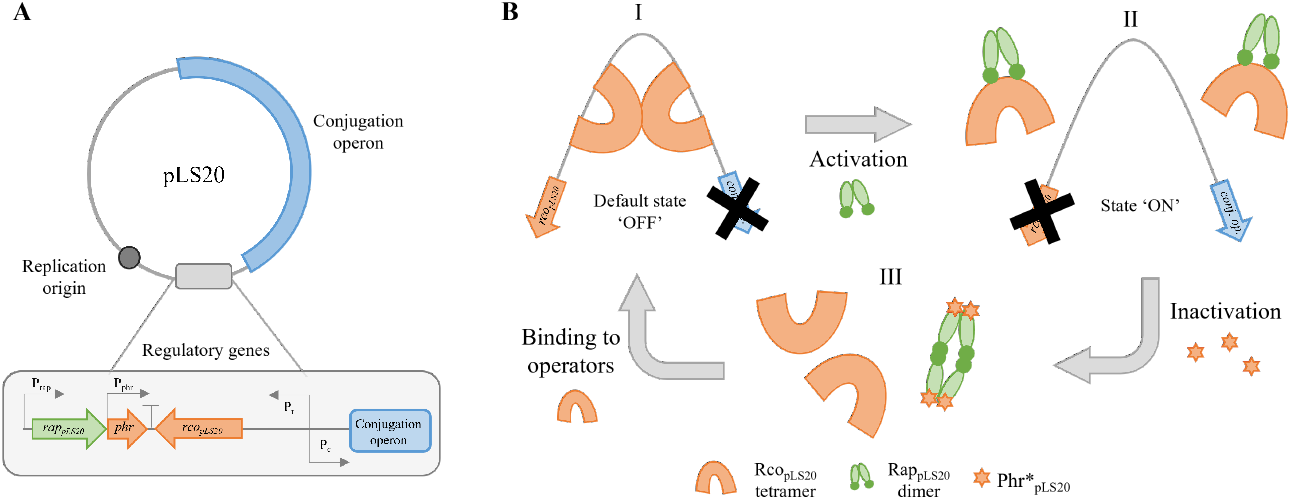
pLS20 conjugative regulatory network and molecular dynamics. **A**. Plasmid map. Orange and green arrows correspond to regulatory genes with an overall repressing or activating effect over the conjugation operon, respectively. **B**. Dynamic stages of the regulatory system. I. Rco_pLS20_ tetramers bound to their operators inhibit the expression of the conjugation operon. II. Rco_pLS20_ tetramers are sequestered by Rap_pLS20_ dimers, allowing the expression of the conjugation operon. III. Phr*_pLS20_ quorum sensing peptides bind Rap_pLS20_ dimers, inactivating their anti-repressor function by inducing tetramerization. Rco_pLS20_ tetramers are then released and can bind their operators again.

Expression of the conjugation operon is regulated by a sophisticated molecular network that integrates environmental and population-level cues (for a review see [13]). The conjugation operon is preceded by a strong promoter, Pc. The activity of Pc is regulated by three proteins whose genes are located immediately upstream of the conjugation operon, Fig 1A. Regulation of the Pc promoter is schematically shown in Fig 1B. The tetrameric transcriptional regulator Rco_pLS20_ represses the Pc promoter by binding to two operators located in the intergenic region between *rco* and gene 28, the first gene of the conjugation operon.

The anti-repressor Rap_pLS20_ antagonizes Rco_pLS20_, relieving repression and activating conjugation. This regulatory cascade is further modulated by the peptide Phr*_pLS20_. Phr*_pLS20_ is a quorum-sensing peptide, produced by donor cells carrying the pLS20 plasmid. It serves as a regulatory signal that controls conjugation activity in response to cell density. Upon binding to Rap_pLS20_, Phr*_pLS20_ inhibits the anti-repressor activity of Rap_pLS20_ under high cell-density conditions, returning the system to its OFF state [14–17], Fig 1B.

The pLS20 regulatory network exemplifies the complexity of conjugation control, where spatial and regulatory dynamics intersect to determine gene transfer efficiency. The interplay of space and regulation on microbial system has been a common topic of computational modeling [18–21]. However, the impact of these dynamics on conjugation at the population level remains poorly understood. Computational approaches, such as individual-based models (IBMs), offer a powerful means to bridge the gap between molecular mechanisms and emergent population behaviors [22–25]. IBMs simulate the interactions of individual cells within their environment, capturing the heterogeneity and spatial structure of bacterial populations. Nowadays, there are several IBM suites, for example: Gro [26, 27], Simbiotics [28], iDynoMICS [25, 29], BSim [30], CellModeller [31, 32], Cell Studio [33] or BioDynaMo [34]. Of these, BSim, CellModeller, and Cell Studio are not optimal for bacterial simulations, since they are designed for the simulation of complex multicellular structures such as biofilms, plant tissues, or immunology phenomena, being computationally costly.

In this work, we use the IBM Gro [26, 27] to investigate the spatial and regulatory factors that influence pLS20 conjugation on solid surfaces. Our simulations reveal how population mixing, colony growth dynamics, and quorum-sensing regulation collectively shape plasmid dissemination. We also develop a complementary Ordinary Differential Equation (ODE) model to capture the macroscopic trends observed in the simulations, providing a broader perspective on conjugation dynamics. Together, these approaches offer new insights into the interplay of spatial organization and regulatory mechanisms in bacterial conjugation, with implications for understanding and controlling HGT in structured environments.

## Materials and methods

### Gro model implementation

We defined a Gro model for the pLS20 conjugative regulatory network [13] as shown in Fig 1. The network operates based on Boolean logic, with each component modeled as either ON (active) or OFF (inactive) based on regulatory conditions. The quorum-sensing signal Phr^*^_pLS20_ acts as the initial input to the system. When Phr^*^_pLS20_ is OFF, Rap_pLS20_ is active and will relief repression of the Pc promoter by co_pLS20_. With Rco_pLS20_ OFF, the conjugation operon is activated, enabling conjugative transfer. Conversely, when Phr^*^_pLS20_ is ON, Rap_pLS20_ is inactive, causing Rco_pLS20_ to suppress conjugation by repressing the Pc promoter. This regulatory network thus functions as a sequential switch, with each state toggling the next in response to the presence or absence of Phr^*^_pLS20_:

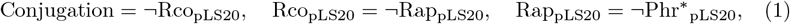

where ¬ represents the Boolean NOT operator. To simplify the model, we excluded self-repression of Rco_pLS20_ at high concentrations [13], as it is irrelevant in the context of a Boolean logic framework. At low concentrations, Rco_pLS20_ activates its own weak promoter, Pr [13]. Therefore, removal of Rco_pLS20_ does not only activate the strong Pc promoter, but also lowers the activity of the already weak Pr promoter, therefore reinforcing the switch-like behavior of the system. Although not explicitly contemplated in the model, this regulation actually aligns with the use of Boolean logic to describe the system.

Within cells, proteins are represented in binary states (active or inactive), whereas the quorum-sensing peptide Phr*_pLS20_ is modeled as a continuous variable in the surrounding medium. Extracellular signal perception is mediated by thresholds that determine when bacterial elements (genes or proteins) shift their binary states. This simplification reflects the biological design of the pLS20 regulatory network, which evolved to operate in a switch-like manner, translating continuous protein concentration dynamics into functional ON-OFF states [13].

Additionally, the model does not include the likely metabolic burden associated with the synthesis of conjugation machinery [35]. This was done because the impact of the burden is not the focus of this work, and, furthermore, to reduce computational cost. The regulatory network is designed to keep the conjugation operon active only in a small proportion of cells, minimizing the burden on the population as a whole.

The implementation of the pLS20 conjugative regulatory network in Gro incorporates the specific parameters listed in Table 1, chosen to model biologically plausible dynamics. In the model, the pLS20 plasmid, described as a binary entity, is only transferable by cell-cell contacts. The expression of its genes into proteins accounts for a delay after gene activation, modeling the time consumed from transcription to translation in living organisms. This delay is defined by the expression time of 1 minute after the gene activation. Once active, virtual proteins present a lifespan defined by the degradation time of 10 minutes. If the degradation timeout is reached and the corresponding coding gene is inactive, the protein state is shifted to inactive (OFF). The quorum sensing peptide Phr*_pLS20_ was included as an extracellular signal molecule secreted continuously from pLS20-containing cells with a rate of 3 min^-1^. Phr*_pLS20_ molecules diffuse with a rate of 0.2 µm^2^/min and degrade with a rate of 0.05 min^-1^. Once the concentration of Phr*_pLS20_ around a cell reaches a threshold of 5 units·cell^-1^, cells uptake the signal at a rate of 2 units·min^-1^, Phr*_pLS20_ is then active inside the cells and the conjugation operon expression is repressed.

**Table 1.** Parameters of the Gro implementation of the pLS20 conjugative regulatory network.

| Parameter (Gro variable) | Value (unit) |
| --- | --- |
| pLS20 conjugation rate (conjRate) | 0.0165 ($\text{min}^{-1}$) |
| Gene expression delay (tacti) | 1 (min) |
| Protein lifespan (tdeg) | 10 (min) |
| $\text{Phr}^*_{\text{pLS20}}$ secretion rate (outFlux) | 3 ($\text{min}^{-1}$) |
| $\text{Phr}^*_{\text{pLS20}}$ diffusion rate (diff) | 0.2 ($\mu\text{m}^2 \cdot \text{min}^{-1}$) |
| $\text{Phr}^*_{\text{pLS20}}$ degradation rate (deg) | 0.05 ($\text{units} \cdot \mu\text{m}^2 \cdot \text{min}^{-1}$) |
| $\text{Phr}^*_{\text{pLS20}}$ import rate (inFlux) | 2 ($\text{units} \cdot \text{min}^{-1}$) |
| $\text{Phr}^*_{\text{pLS20}}$ activation threshold (sensitivity) | 5 ($\text{units} \cdot \text{cell}^{-1}$) |

### Single-cell dynamics of the pLS20 regulatory network

The single-cell behavior of the model was simulated to validate the dynamic properties of the pLS20 regulatory network. The conjugation process transitions between ON and OFF states based on the amount of the quorum-sensing signal Phr*_pLS20_. To evaluate these dynamics, a pulse of Phr*_pLS20_ was introduced between *t* = 10 and *t* = 19 minutes, simulating a transition between quorum and non-quorum states.

Fig 2 illustrates the dynamic shifts of the regulatory components. Initially, both Rco_pLS20_ and Rap_pLS20_ are active, and the conjugation operon is ON. As Phr*_pLS20_ levels increase, Rap_pLS20_ becomes inactive, leading to the activation of Rco_pLS20_ and the repression of the conjugation operon. Upon degradation of Phr*_pLS20_, the system reverts to its initial conjugative active state. The time required for these transitions (approximately 10 minutes) reflects the regulatory network’s responsiveness, allowing cells to adjust their conjugation state within a single doubling period.

**Fig 2.**
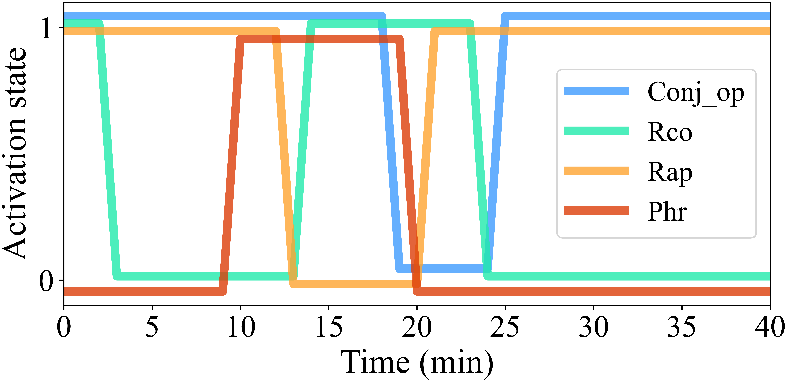
Single cell dynamics of pLS20 conjugative regulatory network. Each line corresponds to a network element. To evaluate the transition dynamics between conjugative states, between *t* = 10 and *t* ≃ 19 minutes, a pulse of the quorum-sensing signal Phr*_pLS20_ is active.

### Gro simulation setup

The Gro simulations were initialized with populations of donor and recipient cells. Donor cells were programmed to secrete Phr*_pLS20_, while recipient cells remained plasmid-free until conjugation occurred. Cell populations were allowed to grow and interact over time, with the spatial dynamics and regulatory network dictating plasmid dissemination. Simulation outputs included cell positions, conjugative states, and the local concentration of Phr*_pLS20_, providing a comprehensive view of pLS20 transmission dynamics.

Donor cells with an active conjugation operon will transfer it to recipient cells to which they are in contact, that therefore become donor cells. For simplicity, we do not contemplate conjugation between donor cells, which is downregulated in any case by a surface exclusion system [36].

Simulation results were averaged over multiple replicates to account for stochastic variability. Error bars in the figures represent standard deviations across replicates. The Gro model implementation and parameter files are publicly available from https://github.com/saulares/pLs20_solid_surface.

## Results

### Impact of Spatial Organization on pLS20 Conjugation Dynamics

The spatial organization of donor and recipient populations plays a critical role in determining the efficiency of pLS20 plasmid conjugation. As an example, Fig 3 shows a simulation starting with an initial donor population placed at the center of a plasmid-free population of recipient cells. Donor cells at the center of the colony accumulate the quorum-sensing signal Phr*_pLS20_, leading to repression of the conjugation operon, while a ring of active conjugation expands the donor population outwards.

**Fig 3.**
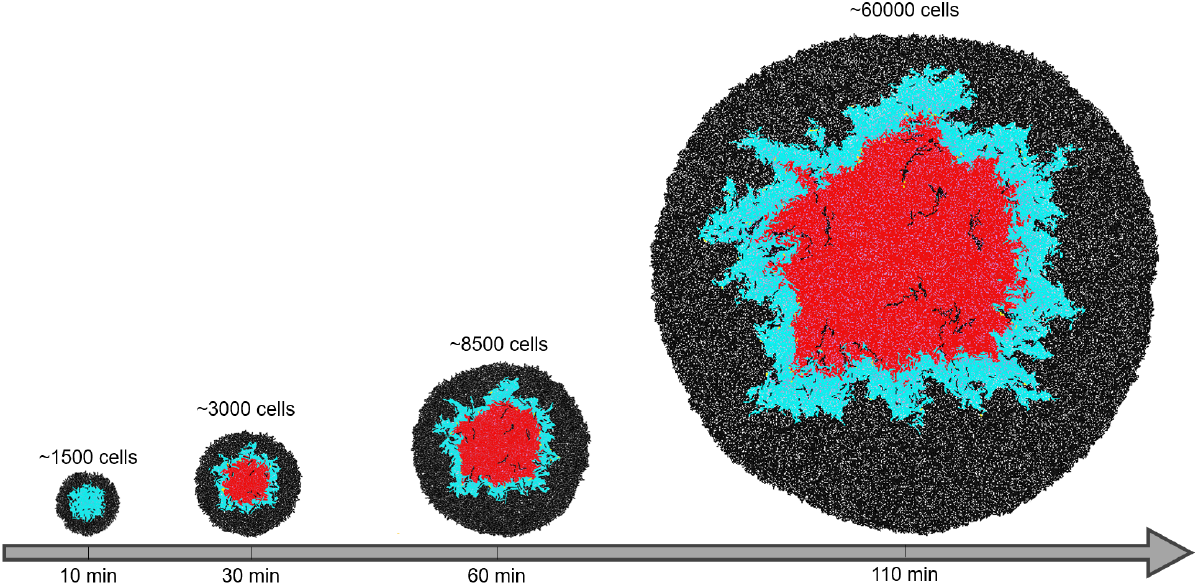
Colony dynamics of pLS20 conjugation. Snapshots of simulations show the spatial spread of the pLS20 plasmid through a colony. The initial population consists of 200 donor cells (blue) at the center of a plasmid-free recipient population (black). Over time, donor cells exhibit repression of the conjugation operon due to Phr*_pLS20_ accumulation (red). Recipient cells trapped within the repressed donor cell regions continue to grow, forming visible filamentous microcolonies at later times.

In this section, we explore the effects of spatial factors such as population mixing and donor placement. Simulations were conducted with varying degrees of donor-recipient mixing, as well as a positive control scenario where donor cells were non-repressible. Fig 4A shows that the efficiency of plasmid transfer is significantly enhanced in highly mixed populations compared to poorly mixed populations, as evidenced by the higher fraction of plasmid-containing cells at the end of the simulation. This is primarily due to the increased contact area between donor and recipient cells in highly mixed populations, which facilitates more frequent conjugative events. For example, the number of plasmid-containing cells at the end of the simulation was 2.5 fold higher in simulations with the highest compared to the lowest degree of mixing (40% versus 16%). This increase emphasizes the importance of spatial proximity and interaction opportunities in determining plasmid dissemination efficiency.

**Fig 4.**
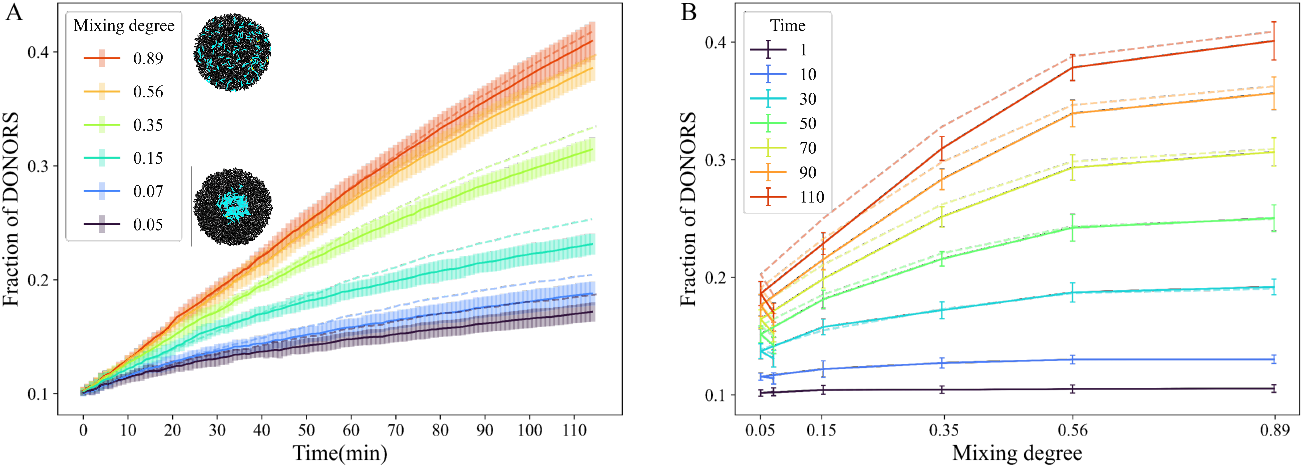
Impact of population mixing on pLS20 conjugation dynamics. Simulations exploring the effects of varying mixing degrees between donor and recipient populations on pLS20 plasmid dissemination. All simulations started with populations containing 10% donor cells. Mixing degree was quantified as the fraction of recipient cells located within the circular area initially occupied by the donor population. **A**. Fraction of donor cells over time for different mixing degrees. Each colored trend represents the mean trajectory from six replicates for a specific mixing degree. Error bars indicate the standard deviation across replicates. Dashed lines correspond to a positive control in which donor cells are non-repressible. Snapshots next to the legend illustrate the initial states of populations with maximum and minimum mixing degrees, color-coded as in Fig 3, where active plasmid-containing cells are blue and recipient cells are black. **B**. Fraction of donor cells for different mixing degrees at various time points. Each trend represents the mean fraction from six replicates for each simulation time, with error bars showing the standard deviation.

The positive control, in which donor cells are non-repressible and thus constitutively active for conjugation, offers an interesting comparison. As shown by the dashed lines in Fig 4, the fraction of plasmid-containing donor cells in this scenario is only marginally higher than in the simulations where repression mechanisms are active. This result suggests that geometric effects, such as the spatial distribution and mixing of cells, can drive conjugation efficiencies close to maximum levels. However, constitutive activity would come at a significant metabolic cost to the donor cells, as they would continuously express and maintain the conjugation machinery. The repression mechanism, therefore, serves as an efficient regulatory strategy to balance conjugation efficiency with metabolic burden, ensuring optimal resource use while maintaining effective plasmid spread.

The local concentration of the quorum-sensing signal Phr*_pLS20_ also plays a significant role in the dynamics of plasmid dissemination. In highly mixed populations, the spatial dispersion of donor cells delays the accumulation of Phr*_pLS20_ reaching threshold levels required to repress conjugation. As a result, donor cells remain active in transferring the plasmid for longer periods of time. In contrast, poorly mixed populations, where donor cells are more clustered, experience higher local concentrations of Phr*_pLS20_, leading to repression of the conjugation machinery and reduced plasmid dissemination efficiency.

Fig 4B corroborates these findings by illustrating how the effects of spatial organization become increasingly pronounced over time. In highly mixed populations, plasmid dissemination continues to increase steadily, driven by sustained interactions between donors and recipients. In contrast, poorly mixed populations exhibit a plateau in donor cell fractions, as quorum-induced repression and limited contact opportunities constrain the spread of the plasmid.

Altogether, this section shows that the degree of mixing between donor and recipient populations is a key determinant of efficient pLS20 plasmid dissemination. While geometric effects alone are sufficient to support efficient plasmid spread, as demonstrated by the positive control, the regulatory repression of the conjugation machinery prevents unnecessary metabolic costs. This highlights the dual importance of spatial organization and regulatory mechanisms in shaping conjugation dynamics, providing valuable insights into the interplay of spatial and metabolic constraints in horizontal gene transfer on solid surfaces.

### Initial colony pattern shapes conjugation efficiency and donor activity

We analyzed how colony growth dynamics on solid surfaces influence pLS20 plasmid transmission by simulating donor populations seeded in concentric rings of varying radii within a recipient cell population. As explained below, these growth-driven spatial redistributions of donor cells were found to significantly affect both plasmid dissemination and the regulation of donor conjugative activity.

Fig 5A demonstrates that the final fraction of donors varies with the initial seeding radius, showing a non-monotonous behavior. Donor populations were seeded in concentric rings of varying radii within a recipient cell population, with the relative radii respect to the initial colony radius ranging from 0.24, close to the center, to 0.88, at the edge of the colony. For the smallest seeding radius, the final fraction of donors is approximately 0.15. As the radius increases, the fraction rises, peaking at 0.36 for radii of 0.63 and 0.73. Interestingly, at a larger radius of 0.88, the final donor fraction decreased to 0.28. This pattern can be attributed to geometric effects inherent in the ring configuration. At very large radii, donor cells are located closer to the colony edge, where they interact with recipient cells on only one side. This limited interaction reduces the efficiency of plasmid dissemination. This finding highlights the intricate relationship between spatial positioning and conjugation efficiency, with donor placement playing a critical role in determining transmission outcomes.

**Fig 5.**
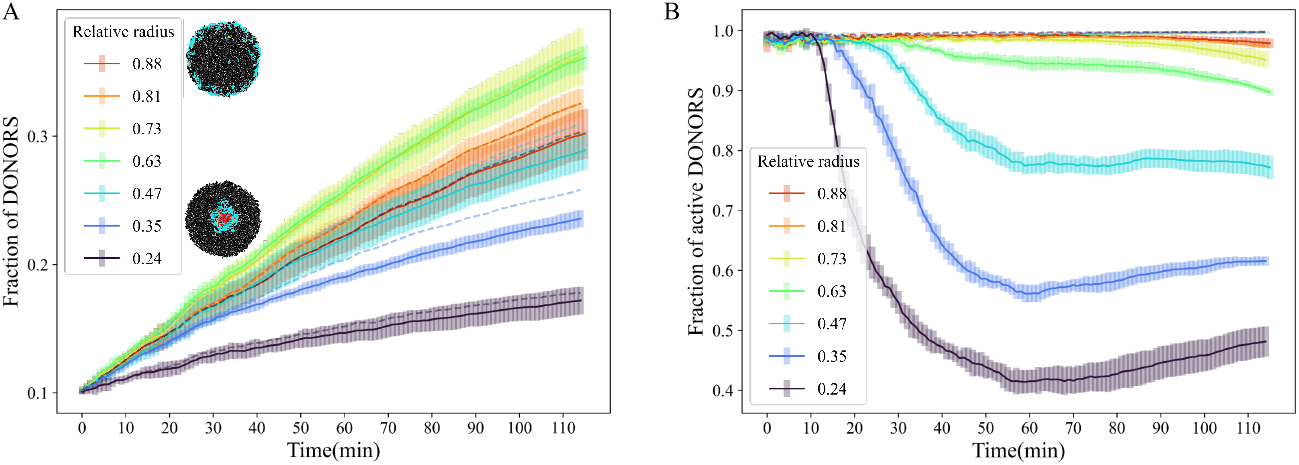
Effect of colony growth dynamics on donor conjugative activity. Simulations of pLS20 plasmid transmission were conducted by seeding donor populations in concentric rings of varying radii within a recipient cell population. All simulations started with 10% donor cells. In both panels, dashed lines correspond to a positive control of non-repressible donors. **A**. Spreading of the plasmid over time. Each trend represents the mean trajectory of 6 replicates for each ring radius. Error bars correspond to the standard deviation among these 6 measures. Snapshots next to the legend represent the initial state of populations with the maximum and minimum seeding ring radius, color-coded as in Fig 3, where active plasmid-containing cells are blue, plasmid-containing cells with repressed conjugation operon are red, and recipient cells are black. **B**. Fraction of donor cells with an active conjugation operon over the total donor population as a function of time. Each colored trend represents the mean fraction of conjugatively active donor cells, relative to the total number of donors, from six replicates for a given seeding radius. Error bars show the standard deviation among replicates.

Fig 5B complements these findings by showing the fraction of donor cells with an active conjugation operon over the total donor cell population as a function of time. Donors seeded at larger radii exhibit higher conjugative activity over time compared to those seeded closer to the center. This difference arises because larger radii promote greater displacement of donor cells during colony growth, reducing the accumulation of Phr*_pLS20_ around them. Conversely, at smaller radii, donor cells remain closer to the center, where Phr*_pLS20_ accumulates to higher concentrations, leading to earlier repression of the conjugation operon. The reduction in conjugative activity at smaller radii occurs sooner and more sharply than at larger radii, as indicated by the transition to the equilibrium phase in Fig 5B.

These results shed light on two aspects of the conjugation dynamics. First, the non-monotonous relationship between the seeding radius and the final donor fraction underscores the impact of spatial geometry on plasmid dissemination. While optimal transmission efficiencies are obtained when donor cells occupy intermediate radii, large and small radii result in suboptimal transmission efficiencies for different reasons. Although the conjugation operon is active due to diffusion of the Phr*_pLS20_ peptide when donor cells are placed at a very large radii, the transmission efficiency is not optimal because the donor cells are close to the edge of the colony, reducing their interactions with recipient cells. Transmission efficiency at very small seeding radii is suboptimal because in these cases Phr*_pLS20_ peptide concentrations accumulate rapidly, resulting in inactivation of the conjugation process. Thus, the spatial pattern of donor cell distribution affects transmission activity in different ways; probability of recipient cell interaction and modulation of the regulatory circuitry by altering accumulation versus diffusion dynamics of the quorum sensing peptide. As a result of these parameters, optimal plasmid transmission efficiencies are obtained at intermediate radii, where donor cells have the highest possibilities to interact with recipient cells and colony growth contributes in mitigating quorum sensing repression.

### An ODE model recapitulates donor population expansion

Agent-based simulations provide detailed insights into the microscopic dynamics of pLS20 plasmid conjugation at the single-cell level. However, Ordinary Differential Equation (ODE) models are often more practical for gaining broader insights into population-level behavior. To complement the agent based simulations, we developed an ODE model describing the dynamics of donor (*D*) and recipient (*R*) populations in a colony where an initial group of donor cells is centrally located and expands outward over time, as shown in Fig 3.

The ODE model tracks the populations of donor and recipient cells with the following equations:

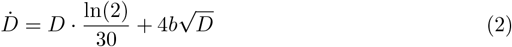

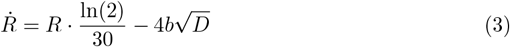

where 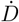 and 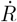 represent the time derivatives of the donor and recipient populations, respectively, and *b* is the conjugation rate parameter.

In the first term in each equation, ln(2)*/*30, corresponds to the growth rate of the bacterial populations. This reflects exponential growth with a doubling time of 30 minutes, which is typical for *B. subtilis* in rich LB medium. Donor and recipient cells are assumed to grow at the same rate, independent of the presence or absence of the plasmid, simplifying the population dynamics.

The second term in the equations accounts for plasmid transmission through conjugation. In this model, conjugation occurs only at the boundary between donor and recipient populations. The number of cells at this donor-recipient interface is proportional to the perimeter of the donor population, which is approximated as 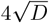, assuming the donor population forms a circular cluster. The conjugation term in 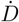 is positive, reflecting the conversion of recipient cells into donor cells, while in 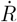, it is negative, representing the loss of recipients as they acquire the plasmid.

The parameter *b* represents the conjugation rate per interaction at the donor-recipient interface. This rate encapsulates the efficiency of plasmid transfer, encompassing molecular and environmental factors that influence the likelihood of a successful conjugation event.

These equations were derived under several simplifying assumptions. The growth of the donor and recipient populations is assumed to be isotropic, with sufficient nutrients and space for colony expansion. Additionally, it is assumed that recipient cells are abundant, ensuring that donor cells always have neighboring recipients to interact with at the colony edge. Conjugation events are limited to a single transfer per donor cell at a time, and conjugated cells immediately transition from recipient to donor without delay. Finally, as in the agent based model, no nutrient or resource constraints are considered. These assumptions simplify the system while maintaining a realistic enough representation of conjugation dynamics.

The resulting ODE model provides a mathematically tractable framework for studying plasmid transmission dynamics. By explicitly linking growth and conjugation terms, the model captures both the biological processes driving plasmid spread and the geometric constraints imposed by spatial organization. This foundation allows for the exploration of how varying parameters, such as initial donor fractions or conjugation rates, influence the overall efficiency of horizontal gene transfer in structured populations.

To validate the ODE model, its predictions were compared with the transmission trajectories obtained from Gro simulations, Fig 6. The fitting process involved adjusting the conjugation rate parameter *b* to align the ODE model with the average trajectories from six independent simulations, each initiated with different donor population fractions. This process yielded a best-fit value of *b* = 0.0165 min^*−*1^, which accurately describes the effective conjugation rate observed in the agent-based simulations. The ODE model showed good agreement with agent simulations for low initial donor fractions. However, for higher donor fractions, deviations became apparent, likely due to the breakdown of assumptions regarding donor-recipient availability and spatial constraints.

**Fig 6.**
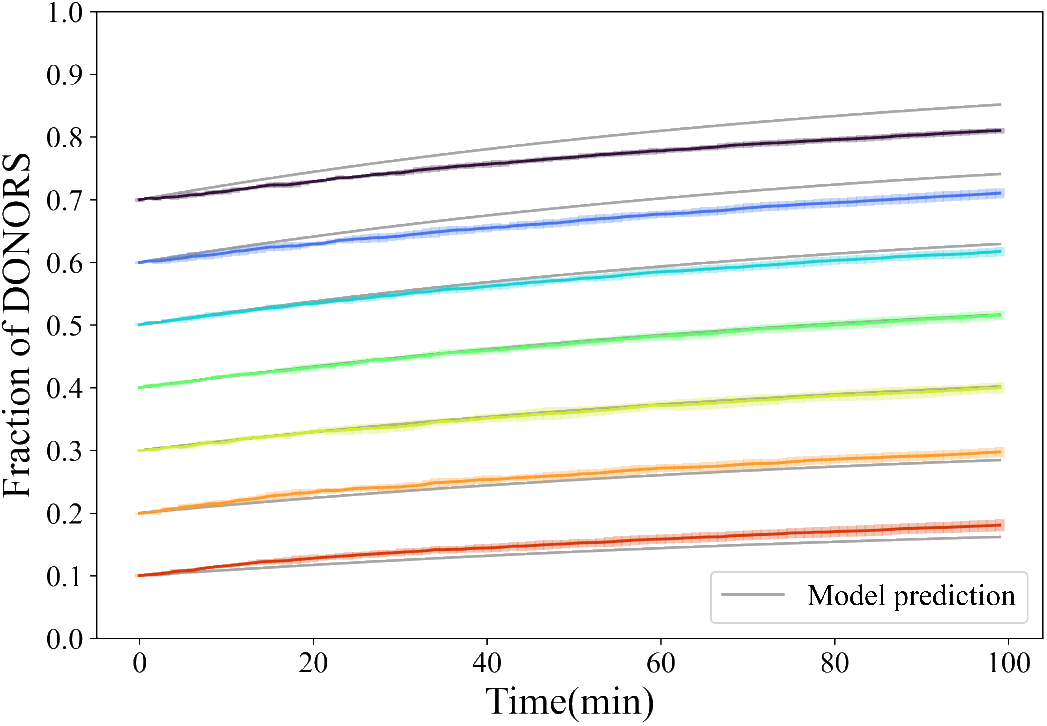
Comparison of ODE model predictions with agent-based simulation results. The colored lines show the mean fraction of pLS20-transformed cells over time from six independent agent-based simulations, with each color corresponding to a different initial donor fraction within the colony. Error bars represent the standard deviation among replicates. Gray lines depict the ODE model fits for each initial donor fraction. A single value of the fit parameter, the conjugation rate *b* = 0.0165 min^*−*1^, is used in all cases.

The comparison between the ODE model and agent simulations provides valuable insights into pLS20 plasmid transmission. While the ODE model effectively captures macroscopic trends in donor expansion and conjugation efficiency, the observed discrepancies for higher donor fractions stress the importance of incorporating spatial dynamics into predictive models. The parameter *b*, representing the conjugation rate, emerges as a critical factor for understanding plasmid transmission on solid surfaces, highlighting the utility of combining agent-based and ODE approaches. This hybrid strategy offers a comprehensive view of both individual-level interactions and population-level dynamics, providing a foundation for further theoretical and experimental exploration of horizontal gene transfer.

## Discussion

This study investigates the spatial dynamics and regulatory mechanisms underlying the conjugation of the pLS20 plasmid in *Bacillus subtilis* populations growing on solid surfaces. Using the agent-based modeling platform Gro and a complementary Ordinary Differential Equation (ODE) model, we provide insights into how colony growth, spatial organization, and quorum-sensing regulation collectively shape plasmid dissemination. Our findings uncover emergent behaviors that reveal the importance of spatial and regulatory interactions in horizontal gene transfer systems.

The spatial distribution of donor and recipient populations was shown to critically influence the efficiency of plasmid transmission. Highly mixed populations achieved significantly higher transmission rates compared to poorly mixed ones, driven by increased donor-recipient contact areas. These results align with the intuitive notion that greater spatial proximity facilitates more frequent conjugative interactions. Importantly, we observed that effective plasmid transmission occurred even under conditions of minimal mixing, while the repression mechanism mediated by Phr*_pLS20_ prevented excessive metabolic burden in the donor cells. This regulatory strategy likely represents an evolutionary optimization that balances the energetic costs of conjugation machinery expression with the need for efficient gene transfer.

Our analysis also revealed the critical role of colony growth dynamics in modulating quorum-sensing regulation. When donor populations were seeded in concentric rings at varying distances from the colony center, donor cells at the periphery of the colony maintained higher conjugative activity compared to those at the center, where Phr*_pLS20_ accumulates rapidly to high concentrations. However, a non-monotonous relationship between the seeding radius and plasmid dissemination efficiency was observed. Intermediate radii resulted in the highest donor fractions, while very large radii exhibited reduced efficiency due to limited interactions with recipient cells at the colony edge. This geometric effect highlights the importance of spatial positioning in determining conjugation outcomes and suggests that donor placement strategies may play a pivotal role in optimizing HGT in structured environments.

The ODE model provided a valuable macroscopic perspective on donor population expansion, capturing the key features of plasmid dissemination while offering a simplified representation of the underlying dynamics. The model accurately described the agent-based simulation results for low initial donor fractions, validating its assumptions under these conditions. However, deviations were observed for higher donor fractions, reflecting the breakdown of model assumptions such as uniform recipient availability. These discrepancies highlight the importance of incorporating spatial heterogeneity into predictive models and emphasize the complementary roles of ODE and agent-based approaches in studying horizontal gene transfer.

Our study also illustrates the potential of agent-based models in bridging single-cell regulatory mechanisms with population-level behaviors. Despite its two-dimensional limitations, the model provided valuable insights into the spatial and regulatory dynamics of bacterial conjugation. Nevertheless, future extensions incorporating three-dimensional colony growth [37, 38], nutrient diffusion [39] and other factors that can affect conjugation could further enhance the realism and applicability of the model.

Overall, our findings underscore the interplay between spatial organization, regulatory mechanisms, and colony growth in shaping the dynamics of plasmid transmission on solid surfaces. The insights gained from this study provide a foundation for exploring strategies to manipulate horizontal gene transfer, with potential applications in controlling the spread of antibiotic resistance and other mobile genetic elements. By combining computational approaches with experimental validation, future work can build on these results to develop more comprehensive models of bacterial conjugation in complex environments.

## Acknowledgments

We thank the support of Elena Núñez-Berrueco and Alfonso Rodríguez-Patón from LIA-UPM. We acknowledge support from grants PID2022-137436NB-I00 (J.B.), PID2022-142185NB-C21 (S.A), PID2022-141969nb-i00 (W.J.J.M), and the research network RED2022-134573-T funded by ‘Ministerio de Ciencia e Innovación’ (MCIU/AEI/10.13039/501100011033) and by ERDF/EU ‘A way of making Europe’. Additional support to J.B. was provided by the E.U. COST action CA22153 ‘European Curvature and Biology Network’ (EuroCurvoBioNet).

MICIU/AEI/10.13039/501100011033 has also funded the ‘Severo Ochoa’ Centers of Excellence to CBMSO, CEX2021-001154-S, and CNB, CEX2023-001386-S.

## References

1. Soucy SM, Huang J, Gogarten JP. Horizontal gene transfer: building the web of life. Nature Reviews Genetics. 2015;16(8):472–482.

2. Daubin V, Szöllősi GJ. Horizontal gene transfer and the history of life. Cold Spring Harbor Perspectives in Biology. 2016;8(4):a018036.

3. Hernando-Amado S, Coque TM, Baquero F, Martínez JL. Defining and combating antibiotic resistance from One Health and Global Health perspectives. Nature Microbiology. 2019;4(9):1432–1442.

4. Salam MA, Al-Amin MY, Salam MT, Pawar JS, Akhter N, Rabaan AA, et al. Antimicrobial resistance: A growing serious threat for global public health. Healthcare. 2023;11(13).

5. Partridge SR, Kwong SM, Firth N, Jensen SO. Mobile genetic elements associated with antimicrobial resistance. Clinical Microbiology Reviews. 2018;31(4):10–1128.

6. Jian Z, Zeng L, Xu T, Sun S, Yan S, Yang L, et al. Antibiotic resistance genes in bacteria: Occurrence, spread, and control. Journal of Basic Microbiology. 2021;61(12):1049–1070.

7. Cabezón E, Ripoll-Rozada J, Peña A, De La Cruz F, Arechaga I. Towards an integrated model of bacterial conjugation. FEMS Microbiology Reviews. 2015;39(1):81–95.

8. Li YG, Hu B, Christie PJ. Biological and structural diversity of type IV secretion systems. Microbiology Spectrum. 2019;7(2):7–2.

9. Singh PK, Meijer WJ. Diverse regulatory circuits for transfer of conjugative elements. FEMS Microbiology Letters. 2014;358(2):119–128.

10. Tanaka T, Koshikawa T. Isolation and characterization of four types of plasmids from Bacillus subtilis (natto). Journal of Bacteriology. 1977;131(2):699–701.

11. Itaya M, Sakaya N, Matsunaga S, Fujita K, Kaneko S. Conjugational transfer kinetics of pLS20 between Bacillus subtilis in liquid medium. Bioscience, Biotechnology, and Biochemistry. 2006;70(3):740–742.

12. Sonenshein AL, Hoch JA, Losick R, et al. Bacillus subtilis and its closest relatives: from genes to cells. Wiley Online Library; 2002.

13. Meijer WJ, Boer DR, Ares S, Alfonso C, Rojo F, Luque-Ortega JR, et al. Multiple layered control of the conjugation process of the Bacillus subtilis plasmid pLS20. Frontiers in Molecular Biosciences. 2021;8:648468.

14. Ramachandran G, Singh PK, Luque-Ortega JR, Yuste L, Alfonso C, Rojo F, et al. A complex genetic switch involving overlapping divergent promoters and DNA looping regulates expression of conjugation genes of a gram-positive plasmid. PLoS Genetics. 2014;10(10):e1004733.

15. Singh PK, Ramachandran G, Ramos-Ruiz R, Peiro-Pastor R, Abia D, Wu LJ, et al. Mobility of the native Bacillus subtilis conjugative plasmid pLS20 is regulated by intercellular signaling. PLoS Genetics. 2013;9(10):e1003892.

16. Singh PK, Serrano E, Ramachandran G, Miguel-Arribas A, Gago-Cordoba C, Val-Calvo J, et al. Reversible regulation of conjugation of Bacillus subtilis plasmid pLS20 by the quorum sensing peptide responsive anti-repressor RappLS20. Nucleic Acids Research. 2020;48(19):10785–10801.

17. Crespo I, Bernardo N, Miguel-Arribas A, Singh PK, Luque-Ortega JR, Alfonso C, et al. Inactivation of the dimeric RappLS20 anti-repressor of the conjugation operon is mediated by peptide-induced tetramerization. Nucleic Acids Research. 2020;48(14):8113–8127.

18. Balagaddé FK, Song H, Ozaki J, Collins CH, Barnet M, Arnold FH, et al. A synthetic Escherichia coli predator–prey ecosystem. Molecular Systems Biology. 2008;4(1):187.

19. Amor DR, Montañez R, Duran-Nebreda S, Solé R. Spatial dynamics of synthetic microbial mutualists and their parasites. PLoS Computational Biology. 2017;13(8):e1005689.

20. Wang T, Weiss A, Aqeel A, Wu F, Lopatkin AJ, David LA, et al. Horizontal gene transfer enables programmable gene stability in synthetic microbiota. Nature Chemical Biology. 2022;18(11):1245–1252.

21. Henderson A, Del Panta A, Schubert OT, Mitri S, van Vliet S. Disentangling the feedback loops driving spatial patterning in microbial communities. npj Biofilms and Microbiomes. 2025;11(1):32.

22. Kitano H. Computational systems biology. Nature. 2002;420(6912):206–210.

23. Hellweger FL, Clegg RJ, Clark JR, Plugge CM, Kreft JU. Advancing microbial sciences by individual-based modelling. Nature Reviews Microbiology. 2016;14(7):461–471.

24. Nagarajan K, Ni C, Lu T. Agent-based modeling of microbial communities. ACS Synthetic Biology. 2022;11(11):3564–3574.

25. Cockx BJ, Foster T, Clegg RJ, Alden K, Arya S, Stekel DJ, et al. Is it selfish to be filamentous in biofilms? Individual-based modeling links microbial growth strategies with morphology using the new and modular iDynoMiCS 2.0. PLoS Computational Biology. 2024;20(2):e1011303.

26. Jang SS, Oishi KT, Egbert RG, Klavins E. Specification and simulation of synthetic multicelled behaviors. ACS Synthetic Biology. 2012;1(8):365–374.

27. Gutiérrez M, Gregorio-Godoy P, Perez del Pulgar G, Muñoz LE, Sáez S, Rodríguez-Patón A. A new improved and extended version of the multicell bacterial simulator gro. ACS Synthetic Biology. 2017;6(8):1496–1508.

28. Naylor J, Fellermann H, Ding Y, Mohammed WK, Jakubovics NS, Mukherjee J, et al. Simbiotics: a multiscale integrative platform for 3D modeling of bacterial populations. ACS Synthetic Biology. 2017;6(7):1194–1210.

29. Lardon LA, Merkey BV, Martins S, Dötsch A, Picioreanu C, Kreft JU, et al. iDynoMiCS: next-generation individual-based modelling of biofilms. Environmental Microbiology. 2011;13(9):2416–2434.

30. Gorochowski TE, Matyjaszkiewicz A, Todd T, Oak N, Kowalska K, Reid S, et al. BSim: an agent-based tool for modeling bacterial populations in systems and synthetic biology. PLoS ONE. 2012;7(8):e42790.

31. Rudge TJ, Steiner PJ, Phillips A, Haseloff J. Computational modeling of synthetic microbial biofilms. ACS Synthetic Biology. 2012;1(8):345–352.

32. Philippou J, Yáñez Feliú G, Rudge TJ. WebCM: A web-based platform for multiuser individual-based modeling of multicellular microbial populations and communities. ACS Synthetic Biology. 2024;13(6):1952–1955.

33. Liberman A, Kario D, Mussel M, Brill J, Buetow K, Efroni S, et al. Cell studio: A platform for interactive, 3D graphical simulation of immunological processes. APL Bioengineering. 2018;2(2).

34. Breitwieser L, Hesam A, de Montigny J, Vavourakis V, Iosif A, Jennings J, et al. BioDynaMo: a modular platform for high-performance agent-based simulation. Bioinformatics. 2022;38(2):453–460.

35. San Millan A, MacLean RC. Fitness costs of plasmids: a limit to plasmid transmission. Microbiology Spectrum. 2017;5(5):10–1128.

36. Gago-Córdoba C, Val-Calvo J, Miguel-Arribas A, Serrano E, Singh PK, Abia D, et al. Surface exclusion revisited: function related to differential expression of the surface exclusion system of Bacillus subtilis plasmid pLS20. Frontiers in Microbiology. 2019;10:1502.

37. Smith WP, Davit Y, Osborne JM, Kim W, Foster KR, Pitt-Francis JM. Cell morphology drives spatial patterning in microbial communities. Proceedings of the National Academy of Sciences of the United States of America. 2017;114(3):E280–E286.

38. Hartmann R, Singh PK, Pearce P, Mok R, Song B, Díaz-Pascual F, et al. Emergence of three-dimensional order and structure in growing biofilms. Nature Physics. 2019;15(3):251–256.

39. Tronnolone H, Tam A, Szenczi Z, Green J, Balasuriya S, Tek EL, et al. Diffusion-limited growth of microbial colonies. Scientific Reports. 2018;8(1):5992.

